# Insights into *in vivo* adipocyte differentiation through cell-specific labeling in zebrafish

**DOI:** 10.1101/2021.03.26.437287

**Authors:** Paola Lepanto, Florencia Levin, Uriel Koziol, Leonel Malacrida, José L. Badano

**Author notes:** shared first authorship (Paola Lepanto, Florencia Levin). To whom correspondence should be addressed: José L. Badano, Paola Lepanto, Institut Pasteur de Montevideo, Mataojo 2020, Montevideo CP11400, Uruguay.

## Abstract

White adipose tissue hyperplasia has been shown to be crucial for handling excess energy in healthy ways. Though adipogenesis mechanisms have been underscored *in vitro*, we lack information on how tissue and systemic factors influence the differentiation of new adipocytes. While this could be studied in zebrafish, adipocyte identification currently relies on neutral lipid labeling, thus precluding access to cells in early stages of differentiation. Here we report the generation and analysis of a zebrafish line with the transgene *fabp4(-2.7):EGFPcaax. In vivo* confocal microscopy of the pancreatic and abdominal visceral depots of transgenic larvae, revealed the presence of labeled mature adipocytes as well as immature cells in earlier stages of differentiation. Through co-labeling for blood vessels, we observed a close interaction of differentiating adipocytes with endothelial cells through cell protrusions. Finally, we implemented hyperspectral imaging and spectral phasor analysis in Nile Red labeled transgenic larvae and revealed the lipid metabolic transition towards neutral lipid accumulation of differentiating adipocytes. Altogether our work presents the characterization of a novel adipocyte-specific label in zebrafish and uncovers previously unknown aspects of *in vivo* adipogenesis.

**Summary statement:** Analysis of the differentiation of adipocytes *in vivo* through cell-specific labeling in zebrafish, revealed their early interaction with blood vessels as well as early lipid metabolic changes.

## Introduction

White adipose tissue (WAT) is present in mammals as well as in the other vertebrates, in the form of anatomically and functionally distinct depots (Zwick et al., 2018). It is formed by adipocytes, adipocyte precursors and macrophages surrounded by a collagen-rich extracellular matrix, and is highly vascularized and innervated. In humans, both visceral and subcutaneous central WAT depots primarily play energy storage and endocrine functions, which are of central importance in the regulation of energy homeostasis. Thus, WAT dysfunction, which is usually associated with obesity, contributes to the development of metabolic syndrome associated-diseases such as type II diabetes, dyslipidemia and non-alcoholic fatty liver disease (Longo et al., 2019). Meanwhile, other depots such as those in the dermis, bone marrow and mammary gland, contribute to regulation of local innate immunity and to the repair of adjacent tissues (Zwick et al., 2018).

In humans, the localization and mode of remodeling of adipose tissue have been associated with healthy or pathological phenotypes (Hepler and Gupta, 2017). Macroscopically, the expansion of subcutaneous WAT (SAT) is considered healthier than the growth of visceral WAT (VAT), as well as its localization in peripheral (extremities) vs central (abdomen in men; abdomen and hips in women) depots. Also, the accumulation of fat can occur in previously existing mature adipocytes (hypertrophy) or in newly differentiated cells (hyperplasia). Importantly, different lines of evidence support an association between adipose tissue hyperplasia with a healthier state as compared to hypertrophy (Vishvanath and Gupta, 2019). Adipogenesis is the process whereby stem cell-like precursors become committed, generating pre-adipocytes which then differentiate into mature adipocytes. While initial formation of adipogenic progenitors occur in hematopoietic tissues (Hudak et al., 2014), adult progenitors reside locally associated to blood vessels of adipose tissue (Hilgendorf et al., 2019; Tang et al., 2008). Tissue environment and cellular composition may influence the differentiation of these locally residing progenitors (for example, (Schwalie et al., 2018)). Thus, the study of adipogenesis and its relationship with other elements in the tissue *in vivo*, is critical to understand normal and pathological processes.

Work on cultured cells has provided key information about transcriptional regulation of adipogenesis (Bahmad et al., 2020). Meanwhile studies in mice have been conducted to analyze the developmental origin of adipocyte progenitors (Hepler and Gupta, 2017). More recently, the use of zebrafish to analyze adipose tissue biology has captured attention as it promises to enable the study of the tissue and its cellular biology *in vivo.* Zebrafish develops only white adipose tissue, which first appears in visceral depots in early larval stages (Flynn et al., 2009; Minchin and Rawls, 2017b). In contrast, mice develop first SAT depots in embryonic stages while VAT appears postnatally (Hudak et al., 2014). Importantly however, zebrafish adipocytes show the same subcellular characteristics, gene expression patterns and final distribution (visceral and subcutaneous) as in mammals (Flynn et al., 2009). Moreover, it has been reported that factors affecting body fat distributions in humans have comparable effects in zebrafish (Loh et al., 2020; Minchin et al., 2015). Thus, taking advantage of its fast external development and optical transparency, zebrafish is an ideal system to study cellular and tisular aspects of WAT development.

Current methods to label adipose tissue *in vivo* rely on the use of lipophilic dyes such as LipidTOX or Nile Red (Minchin and Rawls, 2017a). Nile Red is particularly useful because its absorption and emission spectral characteristics are modified according to the polarity of the environment surrounding the probe (Greenspan and Fowler, 1985). The emission of Nile Red in the context of neutral lipids is blue-shifted in comparison to when it is in the presence of polar lipids. This spectroscopic characteristic has been extensively used to label lipid droplets and to estimate the amount of adipose tissue in live larvae as well as to classify depots (Minchin and Rawls, 2017b). However, Nile Red stains all cell membranes, including the endoplasmic reticulum in which biogenesis of lipid droplets takes place (Olzmann and Carvalho, 2019). Lipid stores are composed of neutral lipids such as triacylglycerols and sterol esters, while polar lipids are present during droplet formation as well as during lipolysis. Maulucci et al. developed an approach to generate a lipid metabolic index using hyperspectral imaging of Nile Red and spectral phasor analysis (Di Giacinto et al., 2018; Maulucci et al., 2018). Thus, this method allowed them to differentiate among cells forced to carry out lipid movement (lipid storage or lipolysis) and those in a resting state. The application of this method to live larvae as well as the study of the interaction of adipocytes with other cells in the tissue would require to specifically label adipocytes independently of their fat load.

To address this we decided to generate an adipocyte specific reporter zebrafish line. Up to date, no factor has been identified that is expressed in all fat depots in mice (Cleal et al., 2017). However, there are several genes that are upregulated in adipocytes in different stages during differentiation, like those used extensively in cell culture of mammalian cells to monitor differentiation progress (Tang and Lane, 2012). As expected, much less information is available from zebrafish. We therefore selected early and late genes which are commonly used as adipocyte differentiation markers in mammalian cell culture models and with previous evidence of being expressed in zebrafish adipose tissue (Imrie and Sadler, 2010), cloned their putative promoter regions and generated transgenesis constructs. We show here that a *fabp4a* −2,7kb proximal genomic region effectively drives the expression of EGFP in adipocytes both previous to and during the accumulation of fat. Membrane tagging of GFP allowed us to observe the early interaction of adipocytes with blood vessels through adipocyte membrane protrusions. Furthermore, we adapted the method of Maulucci et al. by incorporating a three-component analysis in the phasor plot, which enabled us to analyze the lipid metabolism of EGFP-positive cells in live larvae before the formation of lipid droplets. Thus, this new zebrafish transgenic line is a valuable tool which will open new possibilities to study adipocytes and adipose tissue biology *in vivo*.

## Results

### fabp4a(-2.7):EGFPcaax transgene is expressed in early and mature adipocytes

Based on previously reported data on cell culture models and expression patterns in zebrafish we selected four different genes to work with: *adipoqb, cebpa, cfd, fabp4a.* All of them were previously reported to be expressed in zebrafish adipose tissue in larvae and/or adults (Imrie and Sadler, 2010). Taking into account our analysis of the promoter regions, we cloned approximately 2 kb of the proximal part of the promoter for each gene (see Materials and methods), and generated transgenesis constructs using the Tol2 system bearing the cardiac light chain myosin reporter gene, *cmlc2:GFP*, as an early selection marker (Fig. 1A). These constructs were injected together with mRNA coding for Tol2 transposase in the cytoplasm of one-cell stage embryos. Twenty-four hours post-fertilization (hpf) embryos with GFP expressing-cells in their hearts were selected for further breeding. We then analyzed larvae of 15-21 days post-fertilization (dpf) in the stereomicroscope, observing the presence of labelled cells for the constructions with *cebpa* and *fabp4a* promoters. However, only in the latter case the EGFPcaax signal coincided with lipid droplets in mature adipocytes recognizable through transmitted light. Moreover, only in the case of larvae injected with the *fabp4a(-2.7):EGFPcaax* construct we observed mature adipocytes labelled along several generations (Fig. 1B). Therefore, we decided to continue working only with the *fabp4a(-2.7):EGFPcaax* line.

**Figure 1.**
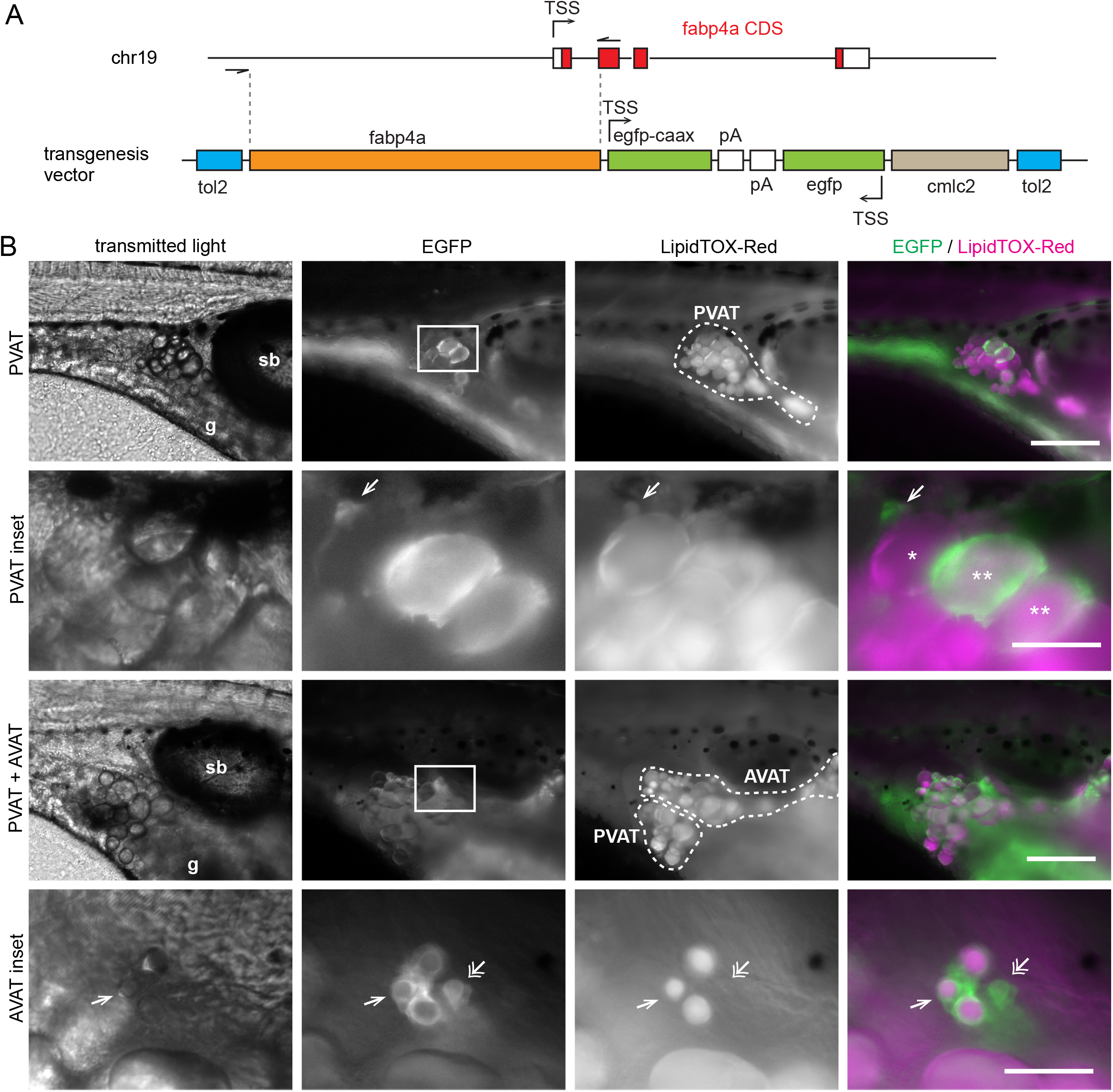
*fabp4a(-2.7):EGFPcaax* is expressed in early and mature adipocytes. **A.** The upper scheme shows the endogenous *fabp4a* gene in chromosome 19 with the transcription start site (TSS), exons (boxes), introns, and the coding sequence (CDS) in red. The cloned region is denoted between dashed lines. The lower scheme represents the vector used for transgenesis (tol2: tol2 sites; pA: SV40_late_polyA; cmlc2: cardiac myosin light chain 2 upstream region). **B.** Epifluorescence microscopy images of live *fabp4a(-2.7):EGFPcaax* larvae from the incross of the F3 generation labelled with LipidTOX-Red. EGFP+ cells were present in the PVAT and AVAT depots. Asterisks indicate mature adipocytes stained with LipidTOX-Red with low (*) or high EGFP expression (**). Single arrows denote early adipocytes expressing EGFP with small lipid droplets. Double arrows indicate early adipocytes with EGFP expression without LipidTOX-Red staining. sb: swim bladder; g: gut. Scale bars: B: panoramic views: 200 μm; insets: 50 μm.

First, to assess EGFP expression in live *fabp4a(-2.7):EGFPcaax* larvae we stained individuals of different stages with the lipophilic dye LipidTOX-Red and analyzed them using epifluorescence microscopy. We observed EGFPcaax signal in the surface of mature adipocytes, both in the pancreatic and abdominal depots (PVAT and AVAT respectively) (Fig. 1B, asterisks). Of note, expression levels varied among cells and this effect remained even after several outcrosses with the wild type fish line. We also analyzed other early-forming depots (renal, ocular and subcutaneous in fins) but failed to detect cells with expression of EGFP. Interestingly, both in PVAT and AVAT, besides mature adipocytes with readily visible lipid droplets, we observed EGFPcaax-positive (EGFP+) cells that had smaller lipid accumulations as well as cells that had no detectable LipidTOX-Red signal (Fig. 1B, single and double arrows respectively). Based on the fact that mammalian *fabp4* is expressed during adipocyte differentiation (Tang and Lane, 2012), these results indicated that EGFP+ cells with no visible lipid droplets (to which we refer as EGFP+/LD- from now on) were likely adipocytes at initial stages of differentiation.

### fabp4a(-2.7):EGFPcaax transgene expression pattern recapitulates the adipose tissue expression domain of endogenous fabp4a

Embryonic *fabp4a* expression has been reported to be restricted to the lens, midbrain and the blood vessels of the head and trunk in 48 hpf embryos (Liu et al., 2007). Meanwhile, 15 dpf larvae present *fabp4a* expression in trunk vessels and in early adipocytes (Flynn et al., 2009). To determine the expression pattern of the *fabp4a(-2.7):EGFPcaax* transgene, we analyzed several developmental stages and compared it with endogenous *fabp4a expression.* For this, endogenous *fabp4a* expression was assessed using fluorescent whole mount *in situ* hybridization (WMISH), and *fabp4a(-2.7):EGFPcaax* transgene expression through immunolabeling with an anti-GFP antibody.

First, we checked the specificities of WMISH probes using *fli1:EGFP* transgenic embryos, which express EGFP in blood vessels, useful as an anatomical reference. For that, 2 dpf wild type embryos were fixed and processed for WMISH as described in the “Materials and methods” section. As specificity controls, we used the *fabp4a* sense probe (negative control) and an antisense probe for *slit2* (additional specificity control). To detect EGFP, we performed a final immunolabeling step. We observed a clear signal corresponding to *fabp4a* transcripts when the WMISH was performed with the antisense probe, which co-localized with *fli1:EGFP* immunodetection almost completely (Fig. S1; note the presence of brain cells positive for *fabp4a* antisense probe without *fli1:EGFP* labeling). Neither of the other two probes generated similar patterns: no specific signal was observed with the *fabp4a* sense probe while the *slit2* antisense probe labeled the ventro-medial part of the neural tube as reported previously (Davison and Zolessi, 2020). These results corroborated the specificity of the *fabp4a* antisense probe and validated the post-WMISH immunofluorescence procedure and reagents.

To compare the distribution of transgenic *fabp4a(-2.7):EGFPcaax* expression with that of endogenous *fabp4a* in larvae, we performed WMISH in individuals of around 21 dpf, immunolabeled them with anti-GFP and analyzed them *in toto* using confocal microscopy. Endogenous expression was observed in the PVAT and AVAT areas with the *fabp4a* antisense probe, coinciding with cells expressing EGFP (Fig. 2A). The specific signal was not observed with the *fabp4a* sense probe (Fig. 2B). Also, EGFP expression was evidenced in pigment cells in live embryos (Fig. S2). This expression pattern was not described before for endogenous *fabp4a* and, in larvae processed for WMISH, expression of endogenous *fabp4a* was not observed in superficial pigment cells (Fig. 2C, single arrows). Instead, we did observe staining for endogenous *fabp4a* in blood vessels (Fig. 2C, double arrows). In conclusion, the expression pattern of the transgene recapitulates the endogenous pattern of *fabp4a* in the adipose tissue, but not within blood vessels or in the brain.

**Figure 2.**
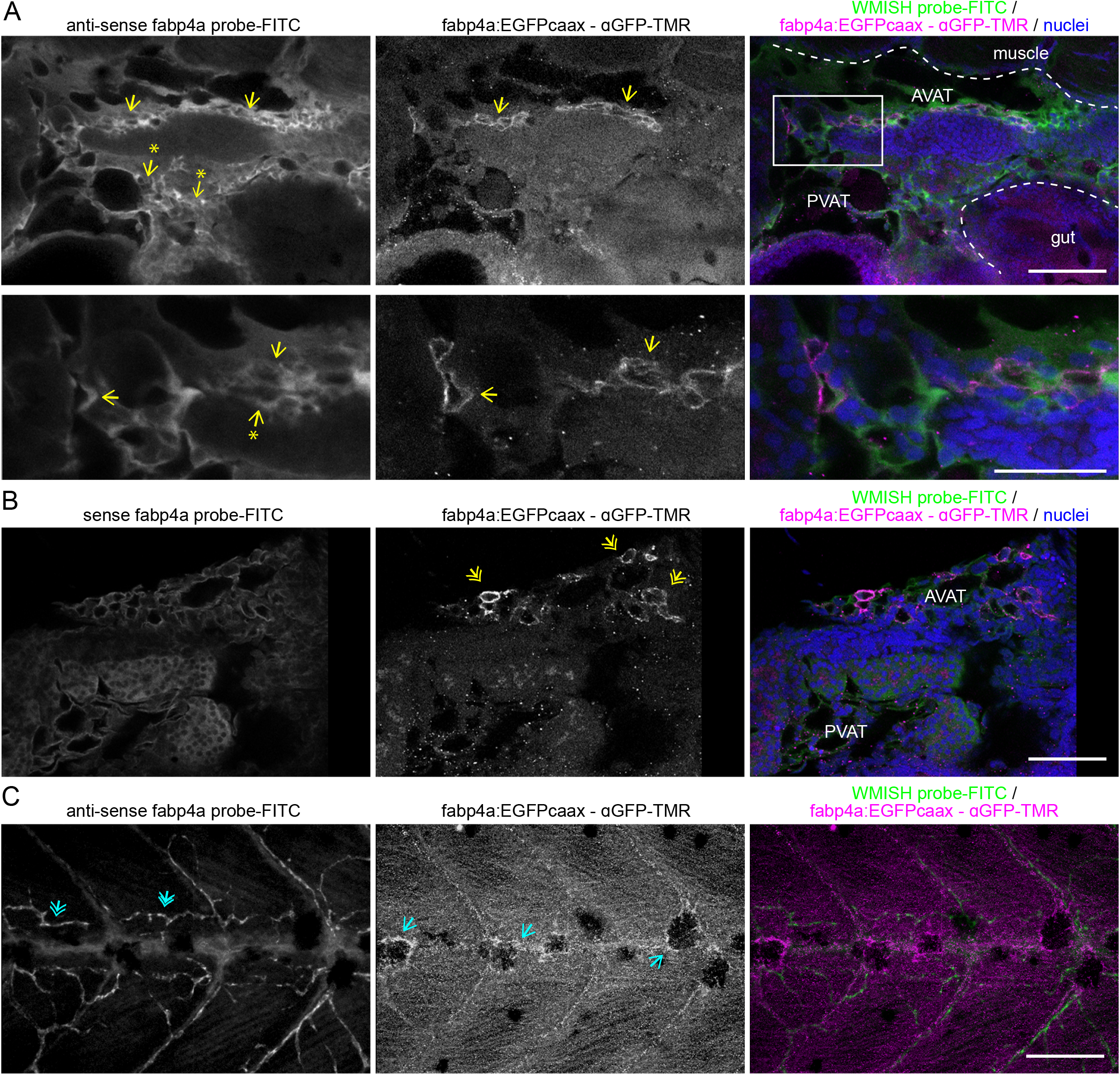
Comparison of the expression pattern of *fabp4a(-2.7):EGFPcaxx* and endogenous *fabp4a* mRNA in larvae. Images of *fabp4a(-2.7):EGFPcaax* larvae of 21 dpf processed for WMISH, immunolabeled with anti-GFP and analyzed *in toto* through confocal microscopy. **A.** Panoramic (upper row) and magnified image (lower row) of the abdominal region of a larvae labeled with *fabp4a* antisense probe. Yellow arrows denote the coincidence of EGFP (immunofluorescence) and WMISH signal. Arrows with asterisks show regions with WMISH labeling and no EGFP signal. **B.** Images of the abdominal region of a larvae labeled with *fabp4a* sense probe. Yellow double arrows indicate regions with EGFP signal without WMISH labeling. **C.** Images of the trunk of a larvae labeled with *fabp4a* antisense probe. Blue single arrows indicate pigment cells with EGFP signal. Blue double arrows show WMISH labeling in blood vessels. Scale bars: A: 100 μm (upper row), 50 μm (lower row); B and C: 100 μm.

### Development of early adipocytes in vivo and their relationship with blood vessels

Our results therefore supported the hypothesis that the *fabp4a(-2.7):EGFPcaax* transgene marks adipocytes during differentiation. To test this we analyzed live larvae and embryos of different stages using confocal microscopy. First, we analyzed embryos in search of early expression of *fabp4a(-2.7):EGFPcaax. In vivo* confocal analysis of 2 dpf and 5 dpf embryos showed expression of EGFP in cells along the antero-posterior axis at dorsal, lateral and ventral positions, all reminiscent of pigment cells (Fig. S2A and C, double arrows). The presence of early labeling of pigment cells suggested that this expression domain corresponded to ectopic expression of *fabp4a(-2.7):EGFPcaax* in the transgenic embryos. To discard autofluorescence or dispersion of light by pigments, we immunostained fixed embryos with anti-GFP antibody. We observed immunostaining co-localizing with EGFP fluorescence (Fig. S2B and D, double arrows), thus confirming that in our fish line pigment cells are labeled by *fabp4a(-2.7):EGFPcaax.*

Previously, it has been reported that lipid droplets are first evident at the right side of the abdomen of early larvae, in ventral and posterior position with respect to the swim bladder (Flynn et al., 2009; Minchin and Rawls, 2017b). Thus we turned our attention into that region in larval stages and stained lipid droplets with LipidTOX-Red. Interestingly, early larvae of standard length (SL) 4.5 mm (8 dpf) show labeling of small cells within the abdomen. At higher magnification we observed the expected surface localization of EGFP. However, these cells did not present lipid droplets (Fig. 3A). We also observed labeling of cells within the trunk in dorsal positions which corresponded to the presence of pigment cells in transmitted light images. Other signals in the images corresponded to autofluorescence of gut contents in the ventral-most part of the larvae and light scattering of pigments in the dorsal half of the swim bladder (Fig. 3A). Despite taking several actions to diminish these confounding signals (16 h food restriction previous to imaging; incubation with epinephrine), they were persistent. Nevertheless, based on the surface localization and intensity of the EGFP signal it was possible to clearly distinguish these EGFP positive and lipid droplet-free cells (EGFP+/LD-).

**Figure 3.**
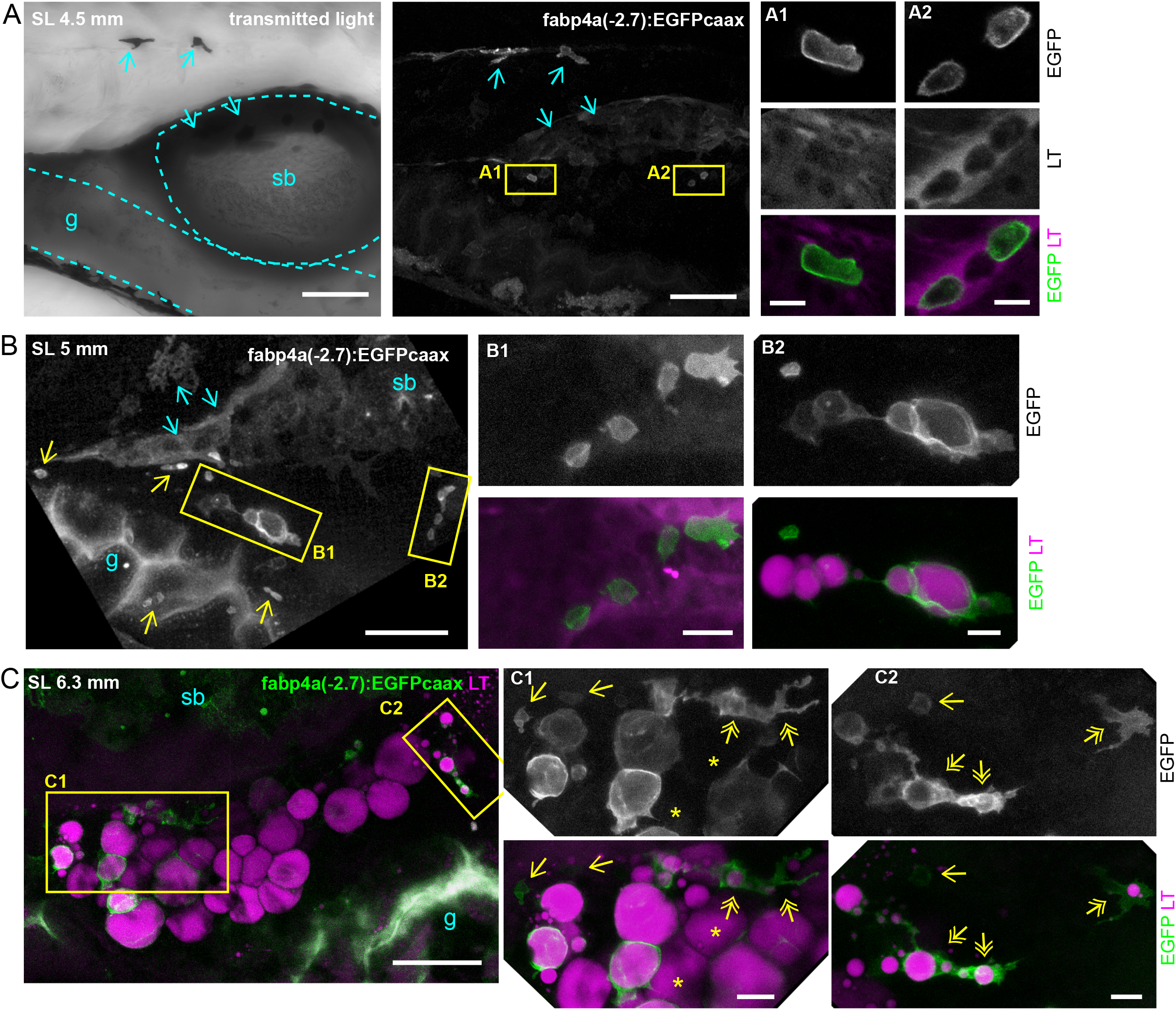
Distribution of labeled cells in the abdominal region of live *fabp4a(-2.7):EGFPcaax* larvae of different stages. Larvae of the indicated stages were stained with LipidTOX-Red, mounted in agarose and imaged using confocal microscopy. **A.** Transmitted light and 3D projection images of a larvae of SL 4.5mm (8 dpf). Yellow rectangles denote cells with transgene labeling. Insets A1 and A2 show confocal sections of these regions. Note membrane localization of EGFP and lack of LipidTOX-Red labeling. Cyan double arrows indicate pigment cells expressing the transgene. Cyan single arrows indicate pigments that scatter light. **B.** 3D projection images of larvae of SL 5 mm (12 dpf). Yellow rectangles denote EGFP+ cells, magnified in B1 and B2. Cells with lipid droplets as well as without them (yellow arrows) can be seen in the same larvae in different positions. **C.** 3D projection images of larvae of SL 6.3 mm (16 dpf) with initial PVAT depot formation. Note the presence of EGFP+ cells with unique cell-filling lipid droplets, irregular cells with several lipid droplets (yellow double arrows) and small cells without lipid droplets (yellow single arrows). Asterisks indicate cells without EGFP expression. Scale bars: A: 100 μm (panoramic view); 10 μm (insets); B: 100 μm (panoramic view); 20 μm (insets); C: 100 μm (panoramic view); 20 μm (insets).

Next, we evaluated older larvae to assess whether these EGFP+ cells in the abdominal region could accumulate lipids. Importantly, larvae of SL 5 and 6.3 mm (12 and 16 dpf respectively) clearly showed cells with surface EGFP signal and lipid droplets of various diameters (Fig. 3B and C). We also noted that some of these cells had irregular forms and projections (Fig. 3C, double arrows). Larvae of these ages also had rounder cells almost completely filled with one big lipid droplet (Fig. 3B and C). In mice, pre-adipocytes have been found to reside near blood vessels (Tang et al., 2008). Thus, we analyzed the relationship of early adipocytes with the vasculature by crossing *fabp4a(-2.7):EGFPcaax* fish with the *kdlr:mCherry* line, which labels endothelial cells. We observed EGFP+ cells both in close apposition and at some distance of vessels (Fig. 4A). Moreover, when cells with lipid accumulation were observed in the PVAT or AVAT depots, some usually appeared in close contact with vessels, sometimes with extensions surrounding them (Fig. 4B).

**Figure 4.**
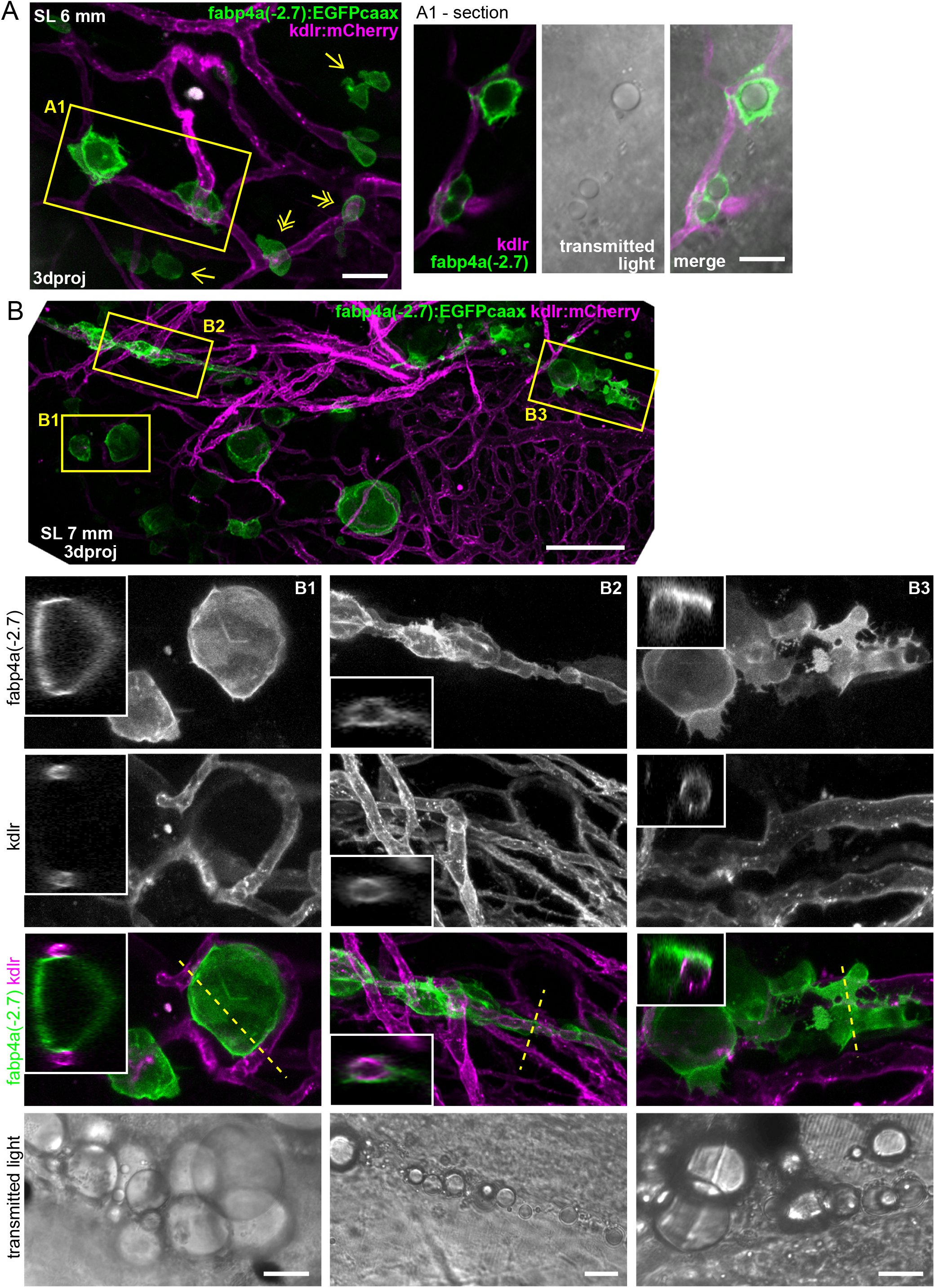
Interaction of early adipocytes with blood vessels. Live larvae from the cross of *fabp4a(-2.7):EGFPcaax* and *kdlr:mCherry* fish lines were imaged through confocal microscopy. Images presented here are 3D projections or single sections, as indicated. **A.** Larvae of SL 6 mm (13 dpf) with many EGFP+ cells in its abdominal area, a few of them having lipid droplets (inset A1). Some of the cells are in contact with blood vessels (double arrows) and some of them are not (single arrows). **B.** Larvae of SL 7 mm (16 dpf), with PVAT and AVAT depots (only some cells of each depot expresses EGFP). Insets B1, B2 and B3 show EGFP+ cells with lipid droplets in close apposition to blood vessels and in some cases surrounding them (B2). 3D projections and sections through the position indicated by the dashed line are shown. Scale bars: A: 100 μm; B: 100 μm (panoramic view), 20 μm (insets).

We expanded our analysis until 21 dpf larvae (larvae of SL 7-8mm) and observed EGFP+ positive cells with different morphologies (Fig. 5A and B). We observed rounded cells filled with a single lipid droplet which likely correspond to mature adipocytes, in some cases having cell projections. Other cells, usually located at the periphery of the depots, typically showed one or more smaller lipid droplets and more irregular morphologies, also with membrane projections (Fig. 5A). In many occasions we observed cells in close proximity to vessels or with extensions surrounding them (Fig. 5B). Remarkably, in all stages analyzed in this work, EGFP+/LD- cells were present. These cells could be observed not only in the abdominal region in AVAT and PVAT but also surrounding the gut in different positions (Fig. 3B), including in the cloaca region (Fig. S3). Interestingly, using transmitted light and high magnification we observed inclusions within EGFP+/LD- cells (Fig. 5C). We performed time lapse acquisitions of those cells for a short period of time. During these time lapse movies we observed that inclusions moved within the cytoplasm (Movie 1). Furthermore, we observed that cells could remain static or have directional movement over a cell diameter distance (Movie 1). This behavior was accompanied by the formation of protrusions which were evident also in our single time point observations (Fig. 3B and 5C).

**Figure 5.**
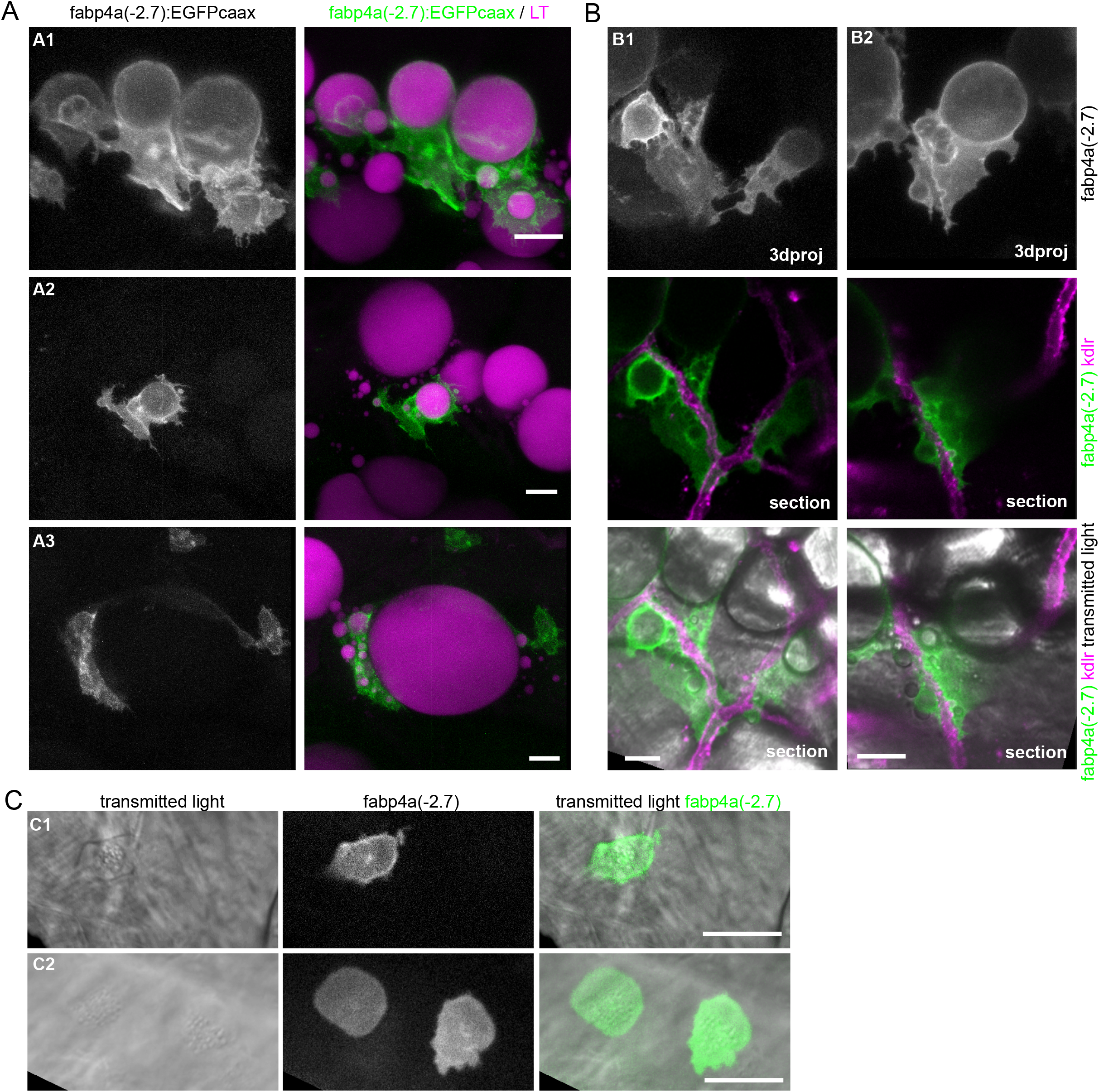
Different cell morphologies observed in *fabp4a(-2.7):EGFPcaax* larvae. **A.** *fabp4a(-2.7):EGFPcaax* larvae of 21 dpf were labeled with LipidTOX-Red and imaged *in vivo* through confocal microscopy. Images are 3D projections of confocal stacks, to show different cell morphologies found in these larvae. **B.** Images of *fabp4a(-2.7):EGFPcaax; kdlr:mCherry* larvae of SL 8 (19 dpf). Note labeled cells in the AVAT depot with cytoplasmic projections which lay in close apposition to blood vessels. Images are 3D projections or sections as indicated. **C.** High magnification confocal sections of EGFP+/LD- cells. Note the cytoplasmic inclusions observed in transmitted light. Scale bars: A: 20 μm; B: 20 μm; C: 20 μm.

Altogether, our results show that *fabp4a(-2.7):EGFPcaax* expressing cells are present in different larval stages, from just before the beginning of the accumulation of fat to later stages where lipid depots are readily visible. Furthermore, we found labeled cells with lipid droplets of different sizes, confirming that cells expressing *fabp4a(-2.7):EGFPcaax* in the abdominal region of larvae are adipocytes in different stages of differentiation. Even though our fish line also expresses EGFP in pigment cells, *in vivo* 3D analysis of the abdominal region efficiently allowed us to distinguish early and mature adipocytes based on localization and cell shape. Furthermore, the results underscore a tight relationship between adipocytes and vessels during their differentiation, and the coexistence of EGFP+ lipid-filled cells with EGFP+/LD- cells in the tissue.

### Analysis of the lipid metabolic profile of early adipocytes with Nile Red fluorescence and spectral phasor plot analysis

As mentioned before, we hypothesized that EGFP+/LD- cells were in fact early adipocytes. Early adipocytes initiate lipid accumulation as part of its differentiation program, and thus would show a mixed lipid environment with neutral and polar components. The quantification of these components has been carried out before in cultured cells through Nile Red fluorescence analysis using spectral phasors (Di Giacinto et al., 2018; Maulucci et al., 2018). In our transgenic larvae, the fluorescence of EGFP could be used as a third component to identify the cells of interest (EGFP+ cells). Thus, we took advantage of the spectral phasor analysis to study the Nile Red spectral shift in the presence of EGFP fluorescence. A similar approach has been used to study membrane polarity using LAURDAN in the presence of mRuby fluorescence (Sameni et al., 2018). For the cellular lipid metabolic profile, wild type larvae stained with Nile Red or *fabp4a(-2.7):EGFPcaax* larvae with or without staining with Nile Red were imaged using hyperspectral detection and the images were analyzed using the advantages of the model-free spectral phasors approach (Fig. S4, the analysis procedure is described in depth in the “Material and methods” section) (Malacrida et al., 2017).

Wild type larvae stained with Nile Red or *fabp4a(-2.7):EGFPcaax* larvae without staining were analyzed first to set the extremes of the distributions in the phasor plot (Fig. S4B). Notice that the Nile Red fluorescence was spread in a trajectory due to the heterogeneity in the polarity of Nile Red environments provided by the intracellular membranes. The position along the trajectory represents pixels with different fractions of membranes with more or less polarity. In the EGFP+ cells labeled with Nile Red, the linear combination for the Nile Red was dragged toward the EGFP position (Fig. S4C). Thus, the extremes of the Nile Red trajectory can be considered as two components and the EGFP as the third component. This strategy enabled us to generate masks for individual EGFP+ cells and to analyze their lipid polarity profile, avoiding the Nile Red signal from other cells. An example image is shown in Fig. S4C. Two cells, one with lipid droplets (cell A) and another without them (cell B), generated clusters at the phasor plot with unequivocally different distribution profiles. To analyze the lipid polarity profile on each of them, we obtained the polarity fractional plot (Fig. S4D). The analysis of cell A yielded a multimodal distribution with higher representation of intermediate zones, whereas cell B gave a single peak in the polar lipid region.

Using this approach, we analyzed EGFP+ cells in larvae at different stages. Representative examples of the observed profiles are shown in Fig. 6A and B. Seemingly mature adipocytes with a big lipid droplet showed a peak in Nile Red profile in regions corresponding to the accumulation of neutral lipids as expected (Fig. 6A and B, “cell D”). Interestingly, it was possible to observe a small peak towards longer wavelengths, representing polar lipid components in the same cells such as the plasma membrane. This was corroborated by the localization of these pixels: the former were localized centrally and the latter surrounded the whole cell (Fig. 6A, see Nile Red profile of “cell D”). EGFP+/LD- cells usually showed distributions enriched in polar components (Fig. 6A and B, “cell A”). Nevertheless, we imaged cells with several peaks or flatter distributions, probably representing transitions between polar and neutral lipid environments (Fig. 6A and B, “cell B” and “cell C”). To summarize and present all the profiles we observed, the center of mass (CM) and distribution range (RD) of the lipid polarity profiles, were calculated and used as characteristics of each distribution for comparison purposes (Materials and methods; Fig. S4D). Within the plot of center of mass vs distribution range (Fig. 6C) it is possible to separate a subgroup of cells with statistically distinct median (for the center of mass and distribution range) and variability (only for the center of mass) compared to those cells outside this region (Fig. 6C and D, dashed line). The low CM and low DR means that the cells within this group are constituted mostly by polar lipids. These cells represent over 50% of the cells analyzed from 8 to 16 dpf (Fig. 6E). The rest of the cells analyzed lay outside the low CM-low DR region due to increasing accumulation of neutral lipids, which extend the DR and bias the CM towards higher values. The percentage of cells analyzed that can be classified in this sub-group increased with larvae age (Fig. 6E). Of note, we observed cells with distinct lipid polarity profiles among larvae of similar standard length and in some cases within the same larvae (Fig. 6E). These observations imply the coexistence of several stages of adipocyte differentiation within the same larvae and suggest that differentiation *in vivo* is continuous and asynchronous. These results indicate that our zebrafish *fabp4a(-2.7):EGFPcaax* transgenic line together with hyperspectral imaging and the spectral phasor analysis shown here is a powerful tool to study changes of the intracellular lipid environment in differentiating adipocytes in live zebrafish larvae.

**Figure 6.**
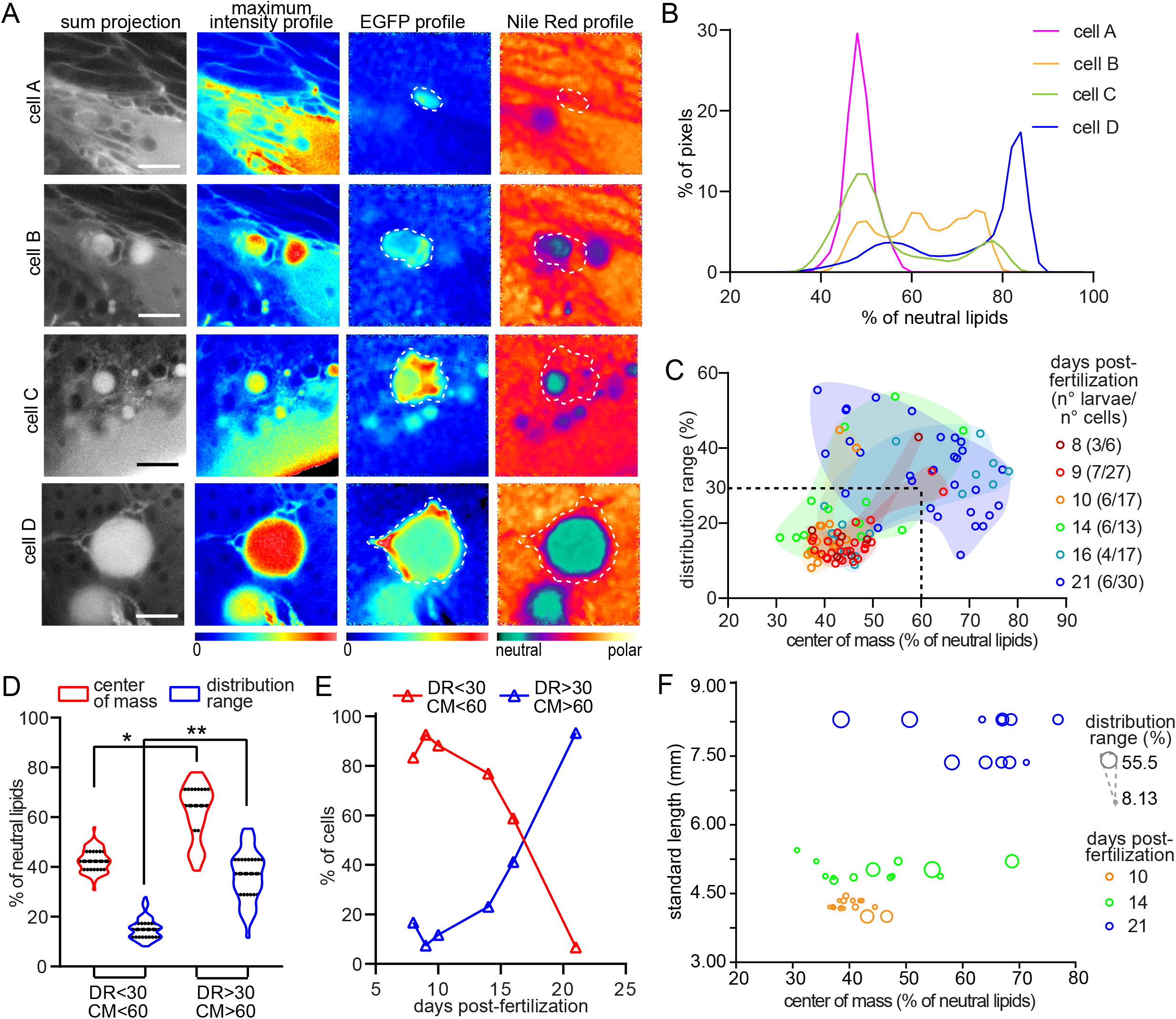
Larvae in different stages present cells with distinct lipid metabolic profiles. **A.** Representative hyperspectral images of adipocytes (“cell A” to “cell D”) in different stages of differentiation. Raw images are presented in gray and intensity based color scale. Images generated after phasor plot analysis make evident the EGFP and Nile Red profiles which are represented separately by different color scales. Scale bars: A: 20 μm. **B.** Plot with the normalized distribution of the image pixels with respect to the fraction (expressed as percentage) of neutral lipids for each cell in (A). **C.** Scatter plot showing the distribution range and center of mass of the cells analyzed in different larval stages. Both variables are expressed as the percentage of neutral lipids. The group of data coming from the same larval stage were enclosed by a manually generated convex hull shape just for visualization purposes. The dashed lines (DR=30; CM=60) generates the two regions considered in (D) and (E). The numbers in brackets indicate the total number of larvae and the total number of cells analyzed in each stage. **D.** Data within (DR<30 / CM<60) and outside (DR>30 / CM>60) the region considered in (C) were separated and compared. For center of mass: (*) p=2.06×10^−13^ for median comparison (Mann-Whitney test), p=1.62×10^−4^ for coefficient of variation comparison (Fligner-Killeen test); (**) p=2.24×10^−16^ for median comparison (Mann-Whitney test), p=7.1×10^−6^ (Fligner-Killeen test). **E.** Representation of the percentage of cells in each group (DR<30 / CM<60 and DR>30 / CM>60) with respect to the larval stage presented in days post-fertilization (dpf). **F.** Representation of the distribution range (as the size of the dots) and center of mass for the cells within some of the larva analyzed; each larva had a different standard length.

## Discussion

A number of studies have started to address the role of compounds on lipid metabolism and storage using zebrafish as a model system (Landgraf et al., 2017). Current knowledge on zebrafish adipose tissue is restricted to fat accumulation capacity using lipophilic dyes due to the lack of specific cell markers. Here we generated and analyzed a new zebrafish line to specifically label adipocytes along their differentiation *in vivo*.

### Expression pattern of fabp4a(-2.7):EGFPcaax

We generated a new transgenic line cloning the proximal part of the promoter of the lipid transporter gene *fabp4a* and used it to direct the expression of a membrane form of EGFP. It was shown previously that *fabp4a* is expressed in the lens, midbrain as well as in blood vessels in the head and trunk of 2 dpf embryos (Liu et al., 2007), an expression pattern that we also observed with our WMISH assay in 2 dpf embryos. However, the *in vivo* analysis of 2 and 5 dpf embryos showed that *fabp4a(-2.7):EGFPcaax* transgene is not expressed in either blood vessels, midbrain or lens cells. In the case of 15 dpf larvae, *fabp4a* mRNA was previously reported to be expressed in abdominal cells with or without neutral lipid accumulation as well as in trunk vessels (Flynn et al., 2009). Our WMISH assay in larvae showed the expression of endogenous *fabp4a* in PVAT and AVAT depots, co-localizing in some cells with expression of *fabp4a(-2.7):EGFPcaax* transgene. Thus, the expression pattern of *fabp4a(-2.7):EGFPcaax* recapitulates primarily the adipose tissue domain of the endogenous expression pattern of *fabp4a*.

We also found that our *fabp4a(-2.7):EGFPcaax* transgene is expressed in surface pigment cells, a domain that does not coincide with the endogenous expression of *fabp4a*. One possible explanation for the lack of expression in some domains as mentioned earlier and the presence of an extra expression domain may be due to the lack of regulatory elements in the cloned region of the promoter. In that case, it would be possible to generate an improved fish line using BAC transgenesis. Alternatively, these extra domains of expression may be due to the action of enhancers present in the genomic region where the transgene was integrated. This is a common drawback of using random insertion of transposons for transgenesis and highlights the importance of using complementary strategies such as insulators or targeted transgene insertion (Caldovic et al., 1999; Roberts et al., 2014). Importantly however, this extra domain of expression did not hinder the utility of the transgenic line as it can be clearly separated from the adipose-related signal by considering the relative position of cells in 3D images.

Another particular aspect of this new transgenic fish line is that the level of expression of EGFP varies among cells. This variability remained even after three outcrosses with the wild type fish line, discarding a mosaicism-based effect. Thus, variation among cells may reflect different cellular states along differentiation which affect transcription from the cloned region of the *fabp4a* promoter, already known to be under a complex regulatory circuit. For example, binding sites for Pparγ and NF-Kβ p50 which modulate transcription in reporter assays, have been reported in the promoter region used in our transgenic line (Laprairie et al., 2017). Moreover, Pparγ has been shown to bind the *fabp4* promoter in a brown adipocyte cell line (Tontonoz et al., 1994). Also, it has been reported that *fabp4* is regulated by VEGFA-DLL4/NOTCH and insulin-FOXO1 pathways in endothelial HUVEC cells (Harjes et al., 2014). In humans, plasma levels of Fabp4, which is mainly produced by adipocytes, have been positively correlated with cardio-vascular disease, type-II diabetes and also with the progression of other diseases by a still undefined mechanism (Prentice et al., 2019). Thus, it would be interesting to analyze which factors contribute to the expression levels of *fabp4* as they may be of clinical relevance, and our results showing significant variability among adipocytes suggests this zebrafish line could be useful to this end.

### Characterization of early adipocytes and their relationship to blood vessels

As mentioned, our detailed microscopic analysis of *fabp4a(-2.7):EGFPcaax* larvae showed labelled cells with different characteristics. In the abdominal region, where WAT depots form, we observed both EGFP+ cells with lipid droplets of various sizes and others without them. These observations are in agreement with previous reports showing that *fabp4a* is expressed in cells with and without lipid droplets in early larvae (Flynn et al., 2009). Accordingly, EGFP+/LD- were present in early larvae (8-10 dpf) well before the initiation of fat accumulation. Notably, we also observed them in older larvae, coexisting with mature adipocytes in fat depots. Our analysis of Nile Red emission of EGFP+ cells/LD- revealed polar lipid or intermediate profiles. Thus, our results indicate that the *fabp4a(-2.7):EGFPcaax* transgene labels adipocytes ranging from early stages of differentiation to mature differentiated cells.

EGFP+/LD- cells presented inclusions when observed with transmitted light at high magnification. These inclusions were LipidTOX-negative and time lapse acquisitions showed that they were highly motile within the cell. Further experiments are required to determine the nature of these inclusions. One interesting possibility is that they may represent initial stages of lipid droplet formation in which the amount of accumulated neutral lipids is not enough to be observed through LipidTOX labeling. LD are formed through accumulation of neutral lipids within the lipid bilayer of the ER, initially forming structures denominated lenses which grow and bud becoming lipid droplets (Olzmann and Carvalho, 2019). Genetic labeling tools that have been developed to evidence initial neutral lipid accumulations may be implemented to study the conservation of early lipid droplet formation mechanisms in zebrafish (Kassan et al., 2013; Wang et al., 2016).

WAT progenitors expressing PPARγ have been reported to reside in the mural compartment of adipose blood vessels in mice (Hilgendorf et al., 2019; Tang et al., 2008). As an analogy to mammals, some authors have hypothesized that WAT progenitors in zebrafish may derive from perivascular pre-adipocytes or, alternatively, from hematopoietic tissue located in the caudal region (Salmerón, 2018). In double labelled larvae, we found EGFP+/LD- cells both in contact and at a distance of blood vessels. In contrast, all EGFP+ cells with lipid droplets were observed in contact with blood vessels. EGFP+/LD- cells were also present surrounding the gut at different positions along the antero-posterior axis. Furthermore, our time lapse acquisitions revealed that these cells had the capacity to migrate. Thus, our results are consistent with the previous formulated hypothesis and *in vivo* time lapse microscopy of EGFP+ cells combined with cell tracing may provide further information. For this, new methods to maintain larvae alive through extended periods of time will be needed, since in our hands, larvae remained alive only for a few hours after mounting in agarose.

Our work also provides information about adipocytes during differentiation and in their mature state. As our transgenic approach included a membrane associated form of EGFP, we could clearly identify the presence of membrane protrusions in early and mature adipocytes. In double-labelled larvae we could appreciate that these membrane protrusions reached blood vessels, suggesting the presence of physical connections. Whether this interaction is direct between cell membranes or indirect through another cell remains to be determined. Extensive evidence supports that several soluble factors coordinate adipogenesis and angiogenesis in obesity as well as in adipose-derived stem cell therapy (Hutchings et al., 2020; Lemoine et al., 2013). Furthermore, secretion of factors by peri-arterial adipocytes can mediate protection or inflammation of the adventitia and atherosclerosis development (Kim et al., 2020). Much less information is available on the interaction of adipocytes and vessels during formation of the adipose tissue (Cao, 2007). Our results suggest an intimate relationship of early adipocytes with blood vessels, probably through cell surface molecules. We hypothesize that these interactions may be instrumental in acquisition of lipids from blood vessels as well as in regulation of growth of the adipose depot.

### Nile Red and phasor approach to characterize in vivo cell lipid metabolism

We used the new *fabp4a(-2.7):EGFPcaax* line to implement a tool for *in vivo* analysis of lipid environment using Nile Red hyperspectral imaging and its analysis through spectral phasor plots (Maulucci et al., 2018). Early adipocytes initiate lipid accumulation as part of their differentiation program, and thus would show a mixed lipid environment with neutral and polar components in their profile. Indeed, as mentioned before, EGFP+/LD- cells showed different profiles, ranging from polar-lipid environment to intermediate polarity-lipid environment. Several groups have studied the lipid composition of *in vitro* differentiating adipocytes of different origins through disruptive methods (Miehle et al., 2020). For example, human undifferentiated adipocytes were enriched in membrane phospholipids such as phosphatidylethanolamines, phosphatidylcholines and sphingomyelins. Meanwhile completely differentiated cells were shown to present diacylglycerols, lysophosphatidylethanolamines and triacylglycerols in addition to membrane phospholipids. Thus our results are consistent with previous analysis, and importantly, provide a base to build on the metabolic analysis of individual cells in their natural context.

Our data show that the technique is sensitive enough to detect lipid environment changes in a non-invasive way and for a specific cell identity, opening the possibility of using this tool to evaluate the progression of differentiation *in vivo* or the effect of drugs on lipid metabolism or genetic interventions. Future development of other fish lines using earlier molecular markers will improve the observation of cells in different stages. For example, work in mice have used *pref1* and *zfp423* to mark adipose tissue progenitors and pre-adipocytes (Gupta et al., 2010; Hudak et al., 2014). Both of these genes are present in zebrafish and may be useful to track the origin of the adipocyte lineage.

### Conclusions and perspectives

In this work we introduced a new zebrafish line labeling adipocytes from early stages up to fully differentiated cells. Furthermore, we described the interaction of early and differentiated adipocytes with blood vessels and evidenced early lipid metabolic changes *in vivo*. We anticipate that the new transgenic line described here will be a useful tool to study the cell biology of adipocytes in the context of the tissue and the whole organism, their interaction with blood vessels and their differentiation *in vivo.* Adipogenesis is highly variable among depots in mammals (Hepler and Gupta, 2017), thus it may be an advantage to use zebrafish for analysis of common conserved cellular mechanisms. Recently, new fish lines labeling lipid droplets have been generated (Lumaquin et al., 2020; Wilson et al., 2021) which may be combined with the *fabp4a(-2.7):EGFPcaax* line presented here for screening approaches focused on genetic and environmental factors affecting early adipocyte differentiation. The *fabp4a(-2.7):EGFPcaax* fish lines and the genetic tools available in zebrafish, combined with two-photon and multiplexing microscopy will surely provide a powerful platform to gain in depth information on adipogenesis and its *in vivo* determinants.

## Materials and methods

### Zebrafish maintenance, breeding and diets

We worked with TAB5 (wild type fish line), *kdlr:mCherry* (blood vessel labeling (Wang et al., 2010)) and *fli1:EGFP* (blood vessel labeling (Lawson and Weinstein, 2002)). *Danio rerio* adults were maintained in a stand-alone system (Tecniplast), at 28 °C, 800 μS/cm^2^, and pH 7.5, with a diet based on live 48 hour-post eclosion *Artemia salina* (artemia cyst from Artemia International) and pellet (TetraMin, tropical flakes, Tetra). Embryos were raised in petri dishes with aquarium water at 28.5 °C (50 larvae per 10 cm petri dish) and bleached at 24 hours post-fertilization. For growth of larvae we used Larval AP100-1 (<50 μm; Zeigler) from 5 to 30 dpf and Golden Pearl Reef & Larval Diet (100-200 μm; Brine Shrimp Direct) from 15 to 30 dpf. Dry food were administered twice per day plus one extra feed of live 24 hour-post hatching *Artemia salina.*

Embryonic staging was performed according to Kimmel (Kimmel et al., 1995) up to 5 dpf and larvae staging (after 5 dpf) was done according to Parichy (Parichy et al., 2009). Standard length (SL) is the distance between the tip of the nose and the caudal peduncle, and it correlates linearly with the growth of adipose tissue as well as the development of other characteristics in larval zebrafish (Minchin and Rawls, 2017b). All protocols (n° 007-19, 009-19, 010-19, 011-19) were approved by the Institut Pasteur de Montevideo ethics committee for the use of animal models (CEUA) and performed by trained, certified staff.

### Promoter cloning

For identification of the potential promoter regions we combined manual analysis and a trial version of Gene2Promoter software (Genomatix). We then designed primers using the Primer-Blast tool from NCBI (Table 1). Candidate primers were blasted against the whole zebrafish genome using the BLAT tool from UCSC Genome Browser. Restriction sites were added at their 5’end to enable directional cloning (underlined in Table 1).

**Table 1.**
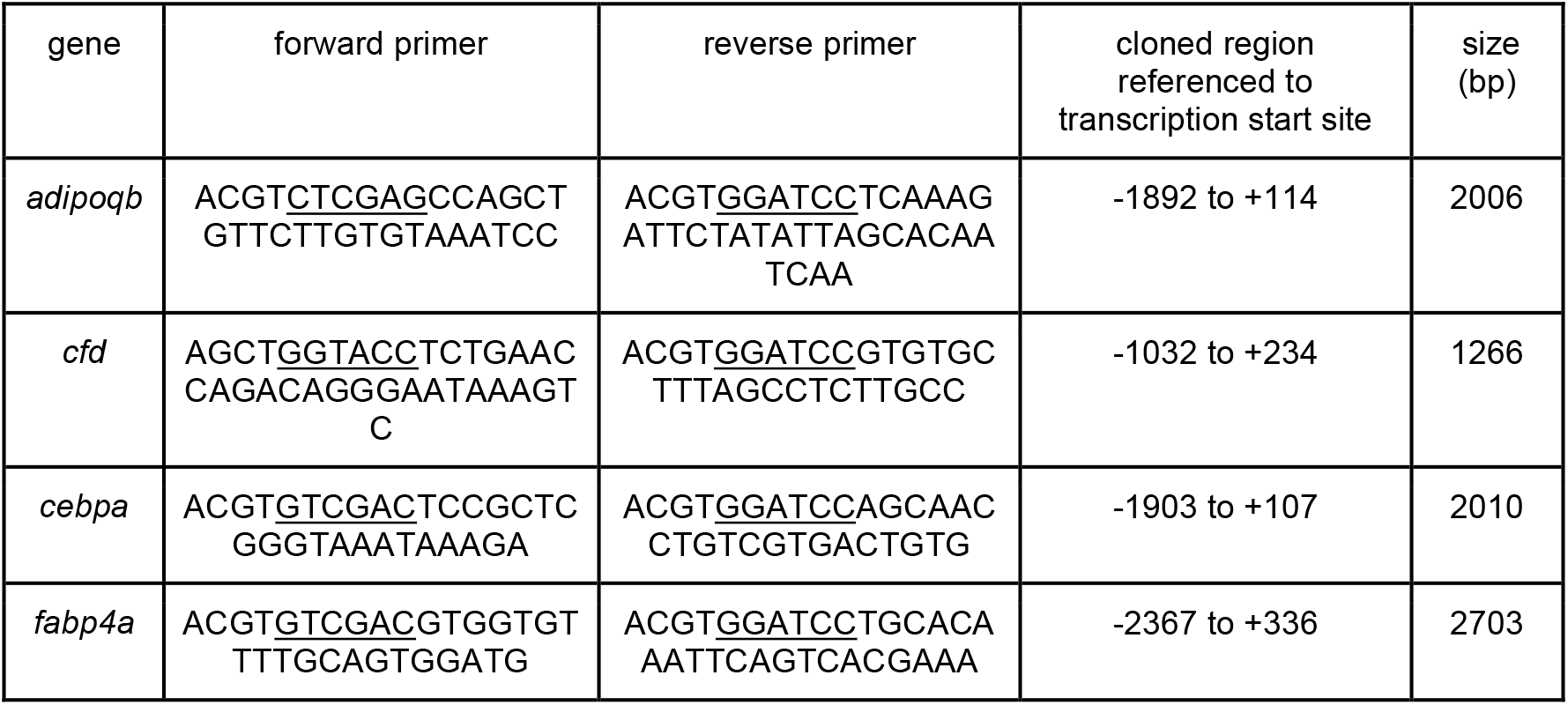
Primers used for amplification of the selected promoter regions, its position in relation to the transcription start site and the size of the amplification product. The underlined regions correspond to the restriction enzyme site.

We used zebrafish high-molecular weight genomic DNA, extracted as previously described (Green and Sambrook, 2014) from 48 hpf TAB embryos. For *fabp4a*, the BAC DKEY-241P5 (Source BioScience) was used as a template. Each region was first cloned into pCRII plasmid using TOPO-TA Cloning Kit (Thermo). After sequencing, they were sub-cloned into p5’Entry-MCS plasmid from Tol2 Kit (Kwan et al., 2007) through digestion and ligation with T4 ligase (Thermo). We recombined each p5’Entry vector with pMiddle Entry vector coding for EGFPcaax (caax is a prenylation signal, directing EGFP to the plasma membrane), p3’Entry Vector with poly-A signal and the pDestTol2-CG2 backbone (with cardiac myosin light chain promoter directing the expression of GFP, *cmlc2:GFP*; and tol2 sites for insertion into the genome).

### Transgenic line generation

TAB5 embryos at one cell stage were injected with 10-20 pg of the desired vector plus 10-20 pg of Tol2 Transposase mRNA. We then selected 24 hpf embryos showing GFP fluorescence in the heart. We grew these embryos until 15-21 dpf when we analyzed the presence of fluorescence in adipose tissue. Selected individuals were outcrossed with the wild-type line until the third generation.

### Fixation, permeabilization and immunolabeling

To decrease pigmentation, embryos were treated with PTU 0.3 % starting at 8 hpf. For the same purpose larvae were anesthetized using tricaine 0.04 g/L, incubated in epinephrine 10 mg/mL plus tricaine 0.04 g/L, mounted in methylcellulose and observed using the stereomicroscope to select larvae expressing GFP. Fixation was carried out in 4 % PFA in PBS overnight at 4 °C.

Fixed embryos and larvae were permeabilized and immunolabeled following the protocol described by Inoue and Wittbrodt with minor modifications (Inoue and Wittbrodt, 2011). Briefly, all steps were carried out at room temperature with agitation unless stated otherwise. Fixed embryos and larvae were washed in PBS plus 1 % Triton X100 (PBST) (3 x 10 mins), dehydrated in a methanol series (50:50 and 100:0 methanol:PBST) (1 x 10 min each) and incubated in 100 % methanol at −20 °C for 20 min. After rehydration in the same methanol series, we performed an antigen retrieval step with 150 mM Tris-HCl pH 9 (5 min at RT and 15 min at 70 °C). After a wash step in PBST (10 min) and two washes in distilled water (5 min each) we further permeabilized samples incubating them in 100 % acetone at −20 °C for 20 min. Finally we washed the samples in PBST several times (6 x 5 min each). For immunofluorescence on WMISH embryos and larvae we followed the same protocol without the acetone permeabilization step.

For immunolabeling all steps were performed with agitation. We incubated permeabilized embryos and larvae in the blocking buffer (10 % FBS plus 1 % BSA in PBST) for 1 h at RT. Primary and secondary antibodies were diluted in the incubation buffer (1 % FBS plus 1 % BSA in PBST). Antibody incubations were performed at 4 °C for 3 days, and washes at RT with PBST. The antibodies used in this work were: anti-GFP (Invitrogen, 1/500), anti-rabbit-633 (Invitrogen, 1/1000).

### Fluorescent Whole-Mount *in situ* Hybridization (WMISH)

We cloned a region of *fabp4a* previously used for probe generation (Flynn et al., 2009) using the following primers: fwd: GATCAAATCTCAATTTACAGCTGTTG; rv: TTCAAAGCACCATAAAGACTGATAAT and oligodT retro-transcribed cDNA as a template. The amplified region was ligated into pGEM T-easy vector (Thermo). Selected clones were checked through digestion and sequencing. The selected clone has the region 195 to 648 from *fabp4a* mRNA sequence, spanning the 3’ half of the CDS and part of the 3’UTR, flanked by T7 and SP6 promoters in 5’ and 3’ respectively. To synthesize the probes we amplified the template using T7 and SP6 primers and afterwards generated digoxigenin (DIG) labeled probes by in vitro transcription with T7 or SP6 polymerases, using Digoxigenin-11-UTP (Merck). As an additional specificity control we used a *slit2* antisense probe which has already been tested (generously provided by C. Davison (Davison and Zolessi, 2020).

The WMISH technique was performed as previously described (Koziol et al., 2014) with modifications following Flynn et al. and Elizondo et al. (Elizondo et al., 2005; Flynn et al., 2009). A detailed protocol is available upon request. Briefly, embryos and larvae were fixed in 4 % PFA prepared in PBS-DEPC water overnight at 4°C. PFA was then replaced twice with 100 % methanol and samples were stored at −20°C until used. After rehydration in an ethanol series, larvae were permeabilized with 15 ug/mL Proteinase K (Fermentas) in PBS-0.1 % Tween-20 (PBS-T) for 10 min (for embryos) or 30 min (for larvae) at room temperature. After a rinse with triethanolamine buffer (0.1 M, pH 8), they were treated twice with acetic anhydride (0.25% v/v for five minutes each), washed with PBS-T, refixed with 4 % PFA in PBS-T for 20 min and washed extensively with PBS-T at room temperature. Pre-hybridization was performed overnight at 60°C in hybridization buffer (50 % formamide, 5X SSC, 1 mg/mL Torula RNA, 100 ug/mL heparin, 1x Denhardt’s solution, 0.1 % Tween-20, 0.1 % CHAPS, DEPC treated water). DIG labelled probes were denatured at 80°C for 3 min and diluted to 0.2 ng/uL in the hybridization buffer. Hybridization was performed at 58°C for 2 days with agitation. Washing steps were done in hybridization buffer at 58°C with agitation, twice for 10 min each, then three times with 2X SSC plus 0.1 % Tween-20 at 58°C for 20 min each, three times with 0.2X SSC plus 0.1% Tween-20 at 58°C for 30 min each, and finally twice with maleic acid buffer (MAB) at room temperature for 15 min. Samples were then blocked overnight at 4°C in 1 % blocking reagent (Roche) plus 5 % sheep serum in MAB and incubated with anti-DIG conjugated to Peroxidase (1/50; Merck) in 1 % blocking reagent diluted in MAB for 3 days at 4°C. Washing steps were done in MAB (three washes of five minutes, followed by three washes of one hour). Fluorophore deposition was carried out with fluorescein-tyramide, prepared and developed as described by Hopman et al. (Hopman et al., 1998). After washes in PBS-T, samples were stored in 80% glycerol in PBS at −20°C until used.

### In vivo labeling and imaging

For *in vivo* lipid labeling, selected larvae were incubated in a 10 cm petri dish (when labeled in group) or 12-well plate (when labeled individually) containing the lipophilic dye diluted in system water. We incubated the larvae with LipidTox Red (Invitrogen, 1/5000) for 1 h at 28 °C or Nile Red (Sigma, 0.78 μM for adipose area quantification and 0.078 μM for emission spectra analysis) for 1 h at 28°C. Labeled individuals were anesthetized and incubated in epinephrine as described above and mounted in 0.8 % low melting point agarose in a 3.5 mm glass bottom petri dish. After solidification, the sample was covered with tricaine 0.04 g/L in system water. To ensure viability during the observation period, a block of agarose covering the region of the gills and the lower jaw was removed using a needle.

*In vivo* images were acquired using epifluorescence or confocal microscopy. For epifluorescence we used an Olympus IX81 with 10x UPlan FLN 0.3 NA and 20x UPlan FLN 0.5 NA Olympus objectives. Confocal microscopy images were acquired with either a Zeiss LSM 800 or Zeiss LSM 880 with a 25x LD LCI Plan-Apochromat 0.8 NA Imm Corr DIC M27 (glycerol, oil, water, silicone) Zeiss objective. Hyperspectral imaging of Nile Red fluorescence was done using the lambda module in the Zeiss LSM 880 with the 25x objective, excitation the 488 Argon laser line was used and the spectra acquisition involved 22 step with 10 nm bandwidth (from 493 nm to 713 nm) using a PMT-GaAsP detector.

### Image analysis

Length and area measurements as well as brightness-contrast adjustments were done using Fiji software (Schindelin et al., 2012).

For Nile Red hyperspectral data analysis we used the spectra phasor approach using Globals for Images SimFCS 4 software (G-SOFT Inc, Champaign, IL-USA). This method transforms the spectral data in each pixel to the real and imaginary component of the Fourier transform, as described earlier by Malacrida et al. (Malacrida et al., 2016):

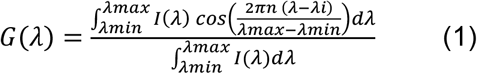

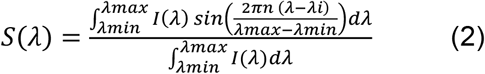

I(*λ*) is the intensity at each step, n the harmonic number and *λ_i_* is the initial wavelength. Each pixel in the image will be located at a single (G, S) position at the spectral phasor plot, yielding a cluster of points due to all pixels in an image. This transformation does not modify the original data and does not involve any fitting or any assumption of components. The position at the phasor depends on the spectrum maximum (phase angle, Θ) and the full width at half maximum (Modulation, M) (Fig. S4A), as:

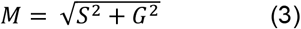

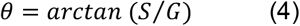

While, red spectral shift implies increasing phase angle, the band narrowing moves the position toward the spectral phasor perimeter (modulation increases).

The spectral phasor plot enables the use of vector properties, such as the linear combination and the reciprocity principle. The linear combination allows the quantification of multiple components in a mixture as a sum fractions of single emitters. In our experiments, Nile Red presented complex photophysics that involved the emission from polar and neutral environments (membrane and lipid droplets, respectively). Furthermore, our phasor plots had an extra component from expression of EGFP. Using the three-component analysis developed by Ranjit and collaborators, we decomposed the fraction of Nile Red in the pixels with EGFP signal (Ranjit et al., 2019). We defined two individual cursor positions (two of the vertices) from the Nile Red trajectory extremes using images from wild-type larvae labelled with Nile Red, and the third position using images of unlabeled *fabp4a(-2.7):EGFPcaax* larvae. The reciprocity principle enables to trace back a region of interest from the spectral phasor (imaginary space) to the original image (real space; the opposite, from a segmentation in the real image to the phasor plot, is also possible). Using this property, we segmented individual cells selecting the corresponding pixels in the phasor plot. Then, we obtained the fractional contributions for the Nile Red trajectory as explained in detail elsewhere (Ranjit et al., 2019). For comparison purposes between different treatments we used the center of mass (CM) for the Nile Red fraction histogram as a central tendency value and the range of the distribution as a dispersion value. The CM for the distribution of each cell Nile Red fraction was calculated as

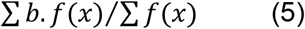

with “*b*” being the percentage of pixels at the particular fraction “*f*(*x*)” of the component “*x*” (Malacrida and Gratton, 2018). The range of the distribution or distribution range (DR) was considered as the *f*(*x*) interval which contains 96% of the pixels. For its calculation we used the accumulated distribution for *f*(*x*) and determined the difference between the *f*(*x*) values corresponding to 2% and 98% of the accumulated distribution.

### Statistical analysis

The statistical analysis was performed using PAST software (Hammer et al., 2001) or Real Statistics Resource Pack software (Release 7.6, Copyright (2013 – 2021), Charles Zaiontz, www.real-statistics.com, accessed on March 2021). For group comparisons we analyzed normality using Shapiro-Wilk test and homogeneity of variances using Levene test. Non-normal and homoscedastic distributions were compared with non-parametric tests (Kruskal-Wallis or Mann-Whitney with Bonferroni correction) as indicated in each case. Non-normal and heteroscedastic samples were rank transformed (Conover and Iman, 1981) and compared using Welch test and Games-Howell post-hoc test. For the comparison of coefficient of variation we used the Fligner-Killeen test. The Reduced Major Axis (RMA) method was used for regression of bivariate data. For comparison of slopes we used the method explained in (Warton et al., 2006).

## Acknowledgements

We thank Luisa Berná for her help defining the putative promoter regions of our genes of interest and Hugo Naya for his advice on statistical analysis. We also thank Gisell Gonzales for her invaluable support in the zebrafish laboratory as well as Magdalena Cardenas and Flavio Zolessi for the revision and helpful comments about the manuscript.

## Competing interests

No competing interests declared.

## Funding

This work was supported by: PEDECIBA and Sistema Nacional de Investigatores-ANII to P.L., U.K., L.M. and J.L.B.; FOCEM - Fondo para la Convergencia Estructural del Mercosur (COF 03/11). LM was supported by the Chan Zuckerberg Initiative.

## Data availability

All the data generated in the study is presented in the manuscript.

## List of Symbols and Abbreviations

hpf: hours post-fertilization
dpf: days post-fertilization
WMISH: whole mount in situ hybridization
PVAT: pancreatic visceral adipose tissue
AVAT: abdominal visceral adipose tissue
SL: standard length.

---

**Figure S1.**
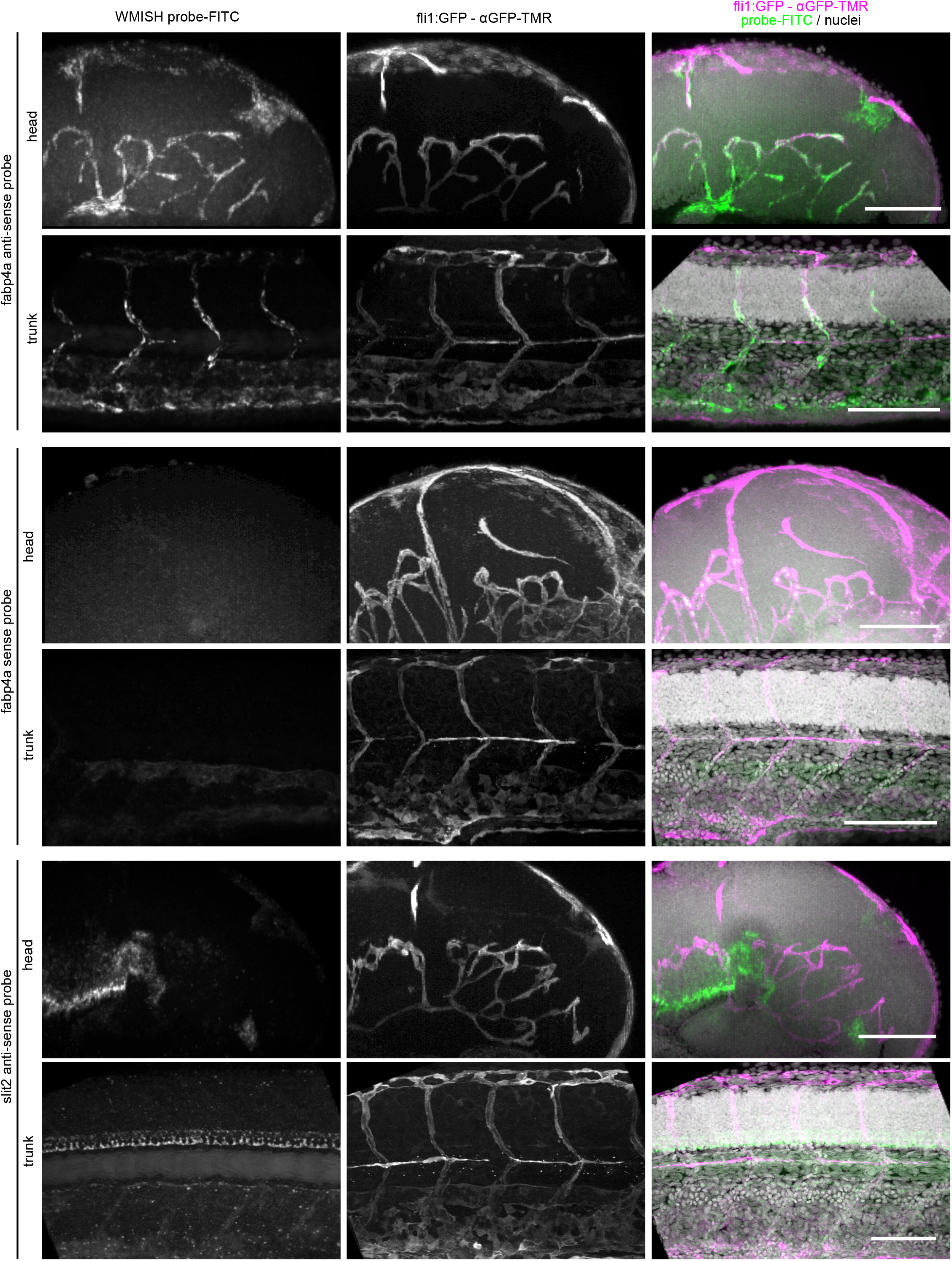
Set up of MWISH and immunofluorescence in embryos. Images of 2 dpf *fli1:EGFP* embryos. WMISH was performed with antisense and sense probes against *fabp4a*, and with previously validated antisense probes for *slit2.* After immunolabeling with anti-GFP, embryos were imaged *in toto* through confocal microscopy. 3D projections generated from confocal stacks are shown. Scale bars: 100 μm.

**Figure S2.**
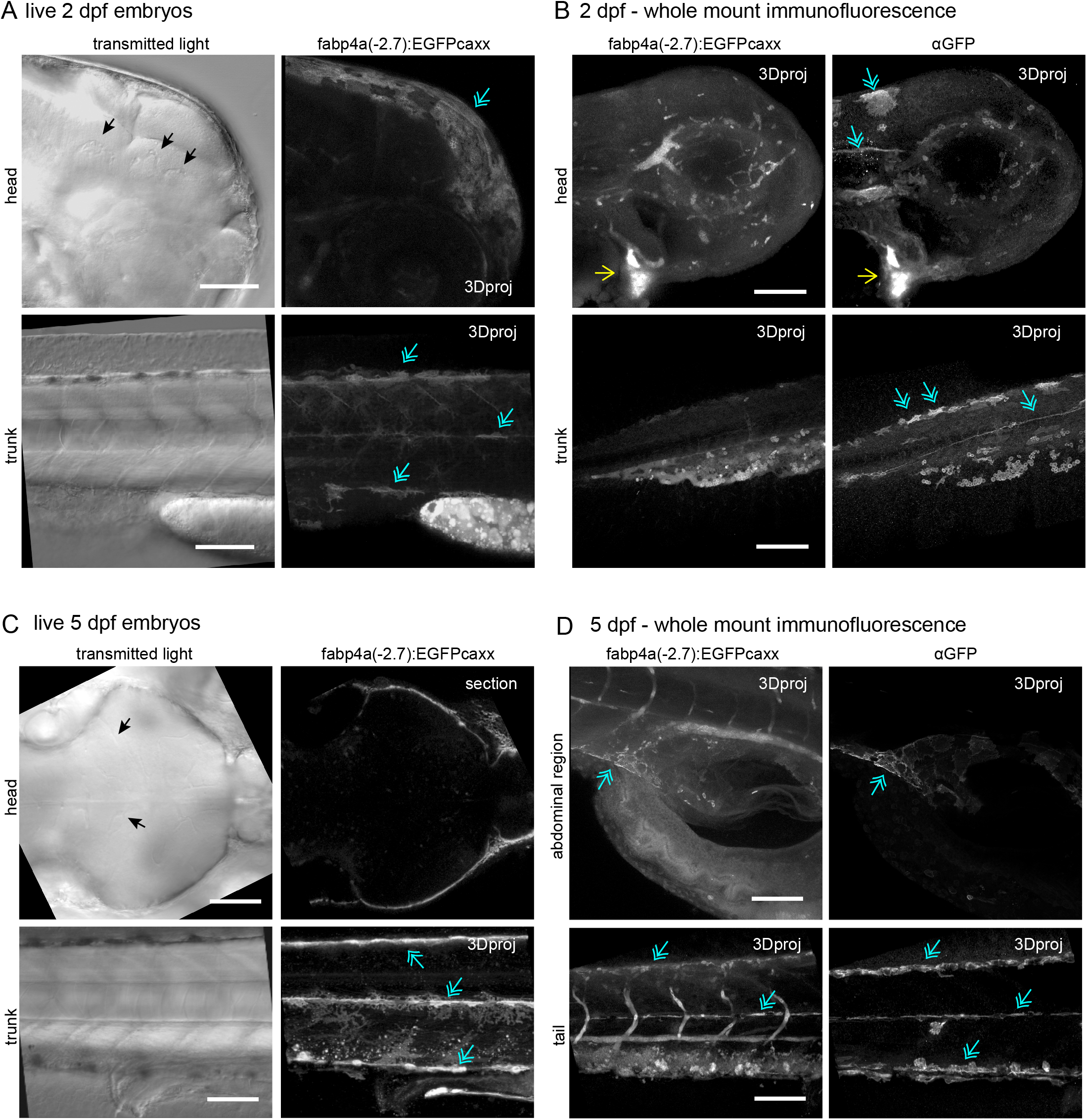
Expression domains of *fabp4a(-2.7):EGFPcaax* in embryos. **A and C.** Images of live embryos of 2 and 5 dpf. Transmitted light or fluorescence images were acquired through confocal microscopy and presented either as single sections or 3D projections (3Dproj). In transmitted light it is possible to observe blood vessels (black arrows) and the lack of fluorescence colocalization. **B and D.** Images of fixed embryos of 2 and 5 dpf immunostained with anti-GFP antibody. Both endogenous GFP and immunolabeling signals are shown. Yellow arrows show the positive immunolabeling signal in heart cells. Blue-double arrows indicate the presence of fluorescence in pigment cells, both in live and fixed embryos, as well as through immunolabeling. Scale bars: A-D: 100 μm.

**Figure S3.**
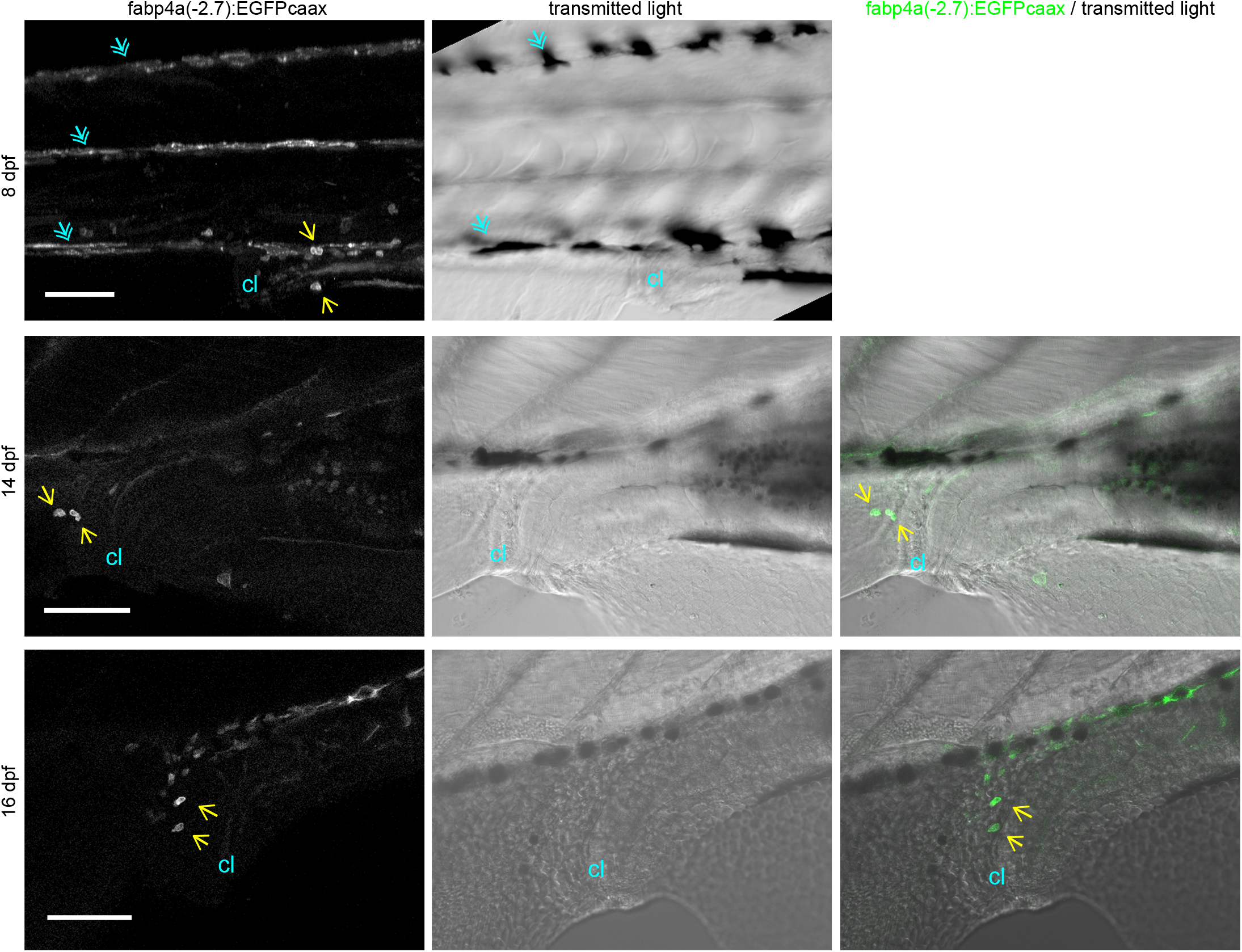
*fabp4a(-2.7):EGFPcaax expressing* cells in the tail region of larvae at different stages. Images of live larvae analyzed through confocal microscopy at different stages (expressed as dpf). EGFP+/LD- cells were present in all stages analyzed, including in the cloaca region (yellow arrows). The fluorescence image of 8 dpf larvae is a 3D projection to note the presence of labeled pigmentary cells (cyan double arrows). cl: cloaca. Scale bars: 100 μm.

**Figure S4.**
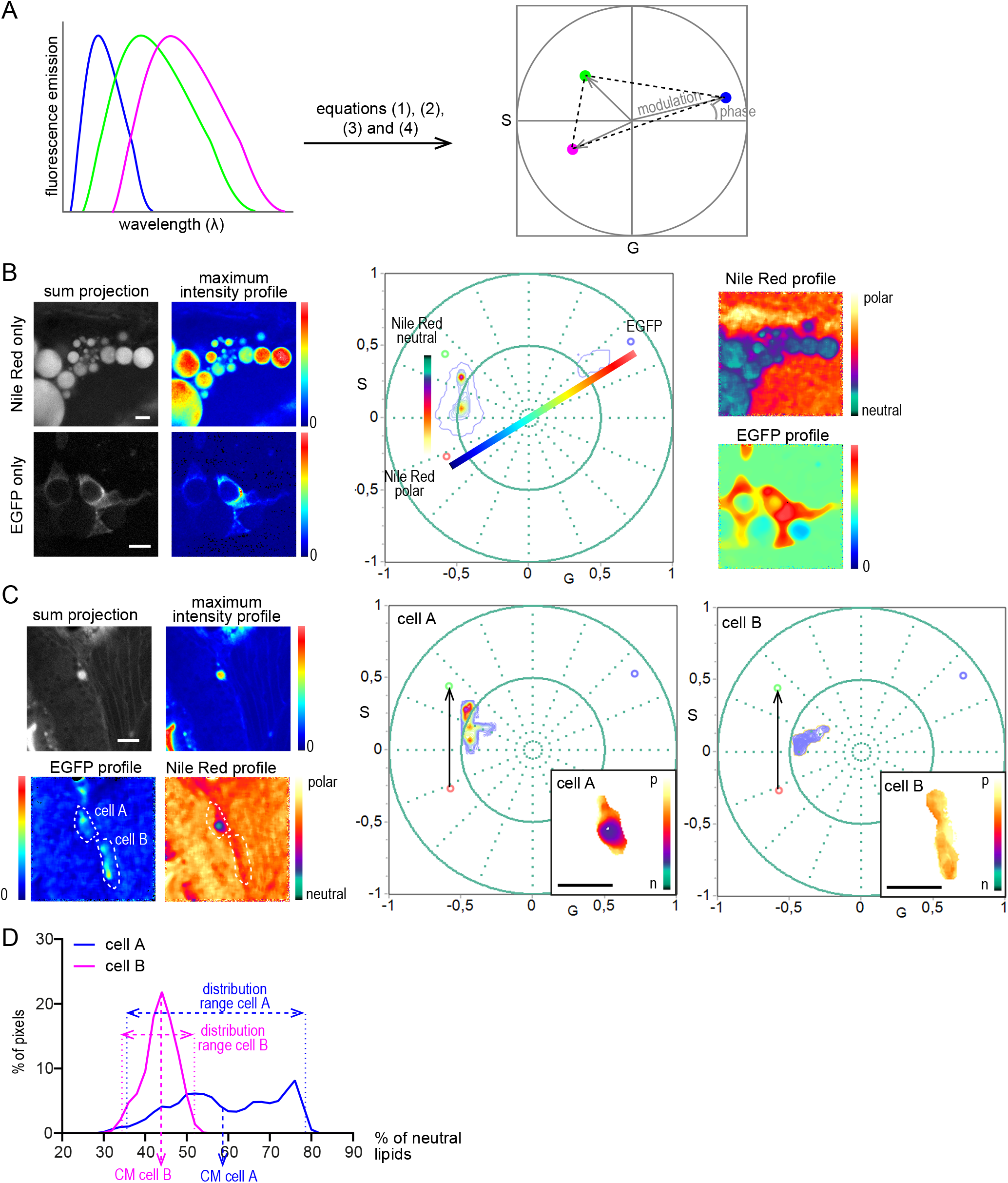
Analysis of the lipid metabolic profile of EGFP+ cells. **A.** Schematic representation of the transformation of the individual spectra of three different fluorophores into the same phasor plot using equations (1) to (4) indicated in “Materials and Methods” section. Dashed lines indicate the area (triangle) in the phasor plot in which pixels with the combination of the three different fluorophores will appear. **B.** Examples of hyperspectral images of control cells and the localization of each pixel in a phasor plot. Images in the left are projections of spectral images and were colored according to pixel intensity maximum. Images of wild type larvae stained with Nile Red (“Nile Red only”) lay within a line-shaped trajectory, corresponding to regions of different lipid polarity within the cell, were used to define the position of two of the components (Nile Red in a polar environment: red circle; Nile Red in a neutral environment: green circle). Instead, images of cells in *fabp4a(-2.7):EGFPcaax* larvae without staining (“EGFP only”) appear in a defined region with low phase angle, which was used to define the position of the third component (EGFP: blue circle). The images in the right were colored according to the position of pixels in the phasor plot (the color scales were superimposed to the Nile Red trajectory and the EGFP axis for improving clarity). **C.** Example of cells in a *fabp4a(-2.7):EGFPcaax* larvae of 8 dpf stained with Nile Red. Images were colored according to pixel intensity or to the position of pixels in the phasor. Phasor plots corresponding to the thresholded cells are presented in the right side. The direction of the Nile Red axis used for plots in (D) is denoted by a black arrow. **D.** Normalized distribution of pixels along the Nile Red axis in the phasor plot for the cells in (C). The “Nile Red axis” corresponds to the percentage of polar lipids in the region of the cell analyzed. The center of mass and range of these distributions were calculated as described in the “Material and methods” section and are schematically represented in the plot. Scale bars: B: 20 μm; C: 50 μm.

**Movie 1.** Time-lapse imaging of EGFP+/LD- cells. The membrane EGFP signal is in green and the transmitted light in grey. Time is showed in minutes:seconds format. Scale bar: 10 μm

## Notes

### Competing Interest Statement

The authors have declared no competing interest.

